# Vascular defects of *DYRK1A* knockouts are ameliorated by modulating calcium signaling in zebrafish

**DOI:** 10.1101/349985

**Authors:** Hyun-Ju Cho, Jae-Geun Lee, Jong-Hwan Kim, Seon-Young Kim, Yang Hoon Huh, Hyo-Jeong Kim, Kyu-Sun Lee, Kweon Yu, Jeong-Soo Lee

## Abstract

*DYRK1A* is a major causative gene in Down syndrome (DS). Reduced incidence of solid tumors and vascular anomalies in DS patients suggests a role of *DYRK1A* in angiogenesis, but *in vivo* evidence is lacking. Here, we used zebrafish *dyrk1aa* mutant embryos to understand *DYRK1A* function in the cerebral vasculature formation. Zebrafish *dyrk1aa* mutants exhibited cerebral hemorrhage and defects in angiogenesis of central arteries in the developing hindbrain. Such phenotypes were rescued by wild-type *dyrk1aa* mRNA, but not by a kinase-dead form, indicating the importance of DYRK1A kinase activity. Chemical screening using a bioactive small molecule library identified a calcium chelator, EGTA, as one of the hits that most robustly rescued the hemorrhage. Vascular defects of mutants were also rescued by independent modulation of calcium signaling by FK506. Furthermore, the transcriptomic analyses supported the alterations of calcium signaling networks in *dyrk1aa* mutants. Together, our results suggest that *dyrk1aa* plays an essential role in angiogenesis and in maintenance of the developing cerebral vasculature via regulation of calcium signaling, which may have therapeutic potential for *DYRK1A*-related vascular diseases.

## INTRODUCTION

The cerebral vasculature plays an essential role in maintaining the homeostasis of the brain by providing oxygen and nutrients and removing wastes. During development, new branches of the cerebral vasculature are formed through angiogenesis by endothelial cells, the primary cell component of the vasculature, via complex cell–cell interactions and signaling pathways from existing vessels (Hogan & Schulte-Merker, 2017). The cerebral vasculature also contributes to the formation of the neurovascular unit (NVU) comprised of pericytes, astrocytes, microglia, and neurons in addition to endothelial cells. Compromise of the normal development or function of the NVU has been implicated in childhood brain development disorders and adult neurological dysfunction (Quaegebeur et al, 2011; Zlokovic, 2008). In the NVU, cerebral endothelial cells, connected mainly by tight junction proteins, build the blood-brain barrier that acts as a primary semipermeable barrier and confers a high selectivity for molecular exchanges between the blood and the brain parenchyma (Dejana et al, 2009). Therefore, the inappropriate development of cerebral endothelial cells may lead to defects in angiogenesis and/or endothelial permeability, which are closely linked to vascular pathologies such as vascular malformations and stroke (Folkman, 1995; Pandya et al, 2006).

The development of the brain vasculature is coordinated by various extra- and intracellular signals. Among them, calcium signaling is one of the major regulators of vascular development and related pathogenesis. Vascular endothelial growth factor signals and various stimuli trigger the change of intracellular Ca^2+^ levels that act as second messengers in endothelial cells, which in turn affect the activity of transcription factors for angiogenesis, such as nuclear factor of activated T-cells, via Ca^2+^-dependent calmodulin/calcineurin activity (Hogan et al, 2003; Loh et al, 1996). In addition, an overload of Ca^2+^ can cause endothelial barrier dysfunction; intracellular Ca^2+^ release via activation of IP_3_ receptors (IP3R) or ryanodine receptors (RyRs) of the endoplasmic reticulum into the cytoplasm can increase vascular permeability through disorganization of VE-cadherin or cytoskeletal rearrangement (Gao et al, 2000; Shen et al, 2010; Tiruppathi et al, 2002). Intriguingly, it has been reported that calcium supplementation for bone health can unexpectedly induce stroke and cardiovascular diseases (Reid & Bolland, 2008; Wang et al, 2014), and calcium channel blockers and calcium antagonists have been clinically used as therapeutic agents for strokes and blood vessel dysfunction (Inzitari & Poggesi, 2005). Thus, understanding the detailed underlying molecular mechanisms of Ca^2+^ signaling in angiogenesis and vascular permeability along with the identification of key players may provide an important therapeutic means for treating vascular diseases.

Dual-specificity tyrosine phosphorylation-regulated kinase 1A (DYRK1A) is a serine-threonine kinase that has a dual kinase activity capable of autophosphorylating its own tyrosine residues and phosphorylating other substrates. The *Dyrk1a* gene was first identified in a *Drosophila* screening as a mutant *minibrain*, whose name came from the brain morphology defects with reduced brain size (Lochhead et al, 2005; Tejedor et al, 1995). *Dyrk1a* is located in a Down syndrome critical region (DSCR) and is best known as a major causative gene implicated in brain function, neurological defects, and neurofibrillary tangle formation in Down syndrome (DS)(Liu et al, 2008; Wegiel et al, 2011). Interestingly, epidemiological studies suggested that DS patients have a reduced incidence of angiogenesis-related solid tumors (Hasle et al, 2000; Nizetic & Groet, 2012) and carry numerous vascular anomalies such as umbilico-portal system anomalies, vertebral and right subclavian artery defects, and pulmonary vein stenosis (Gowda et al, 2014; Pipitone et al, 2003; Rathore & Sreenivasan, 1989; Stewart et al, 1992). Furthermore, they also have an increased incidence of Moyamoya disease and cerebral amyloid angiopathy, which are associated with a cerebrovascular dysfunction and intracerebral hemorrhage (de Borchgrave et al, 2002; Donahue et al, 1998; Jastrzebski et al, 2015; Mito & Becker, 1992; Sabde et al, 2005). Consistent with the potential role of *DYRK1A* in angiogenesis, the TS65Dn mouse model of DS, trisomic for the *Dyrk1a* gene, exhibited reduced tumor growth, presumably by suppressing tumor angiogenesis(Baek et al, 2009). Taken together, DYRK1A may be implicated in vascular formation/function, and this could provide a new prospective to understanding DYRK1A-related pathogenesis; however, *in vivo* evidence of DYRK1A function in vascular pathology is scarce.

To investigate the role of DYRK1A in vascular formation, we adopted developing zebrafish as a model organism. Zebrafish is a vertebrate animal model used for genetic studies of human diseases exhibiting a high similarity to humans at anatomical and molecular levels, especially in the vascular and nervous system (Isogai et al, 2001; Schmidt et al, 2013). Zebrafish embryos can be readily manipulated for genetic gain-of-function studies with transgenesis or mRNA overexpression, and loss-of-function studies with gene knockouts or morpholino use (Clark et al, 2011; Hogan et al, 2008; Timme-Laragy et al, 2012; Varshney et al, 2015; Zu et al, 2013). Large clutch sizes and various inexpensive and fast experimental techniques allow use of zebrafish for unique high-throughput *in vivo* small molecule screening, which enables the identification of hit compounds and gives insights into potential underlying mechanisms(MacRae & Peterson, 2015).

We have recently reported autistic behavioral phenotypes of knockout mutants of *dyrk1aa*, a mammalian *DYRK1A* homolog, named *dyrk1aa*^*krb1*^, generated by transcription activator-like effector nucleases (Kim et al, 2017). In the current study, we investigated a role of *dyrk1aa* in cerebrovascular development during embryogenesis using *dyrk1aa* loss-of-function mutants. The *dyrk1aa*^*krb1*^ mutants exhibited cerebral hemorrhage and angiogenic defects in the developing hindbrain, as analyzed at high resolution by confocal fluorescent microscopy and transmission electron microscopy, using transgenic animals. These vascular abnormalities were rescued by expression of wild-type *dyrk1aa* mRNA, but not a kinase-dead form, indicating an essential role of its kinase activity. Chemical screening using an FDA-approved chemical library identified the calcium chelator ethylene glycol-bis(β-aminoethyl ether)-N,N,N’,N’-tetraacetic acid (EGTA) as efficiently rescuing cerebral hemorrhage as well as abnormal cerebrovascular defects in *dyrk1aa* mutants. Another calcium signaling modulator, FK506, rescued the hemorrhagic and cerebrovascular defects of *dyrk1aa* mutants in a similar manner as EGTA, and transcriptomic analyses identified changes in calcium signaling as the main pathway affected in *dyrk1aa* mutants. Together, the cerebral hemorrhage and cerebrovascular defects of our zebrafish *dyrk1aa* mutants and our chemical screening revealed an important but less-known role of *DYRK1A* in *in vivo* vascular formation, which involves a mechanism mediated by calcium signaling, providing a potential therapeutic target for DYRK1A-related vascular disorders.

## RESULTS

### Cerebral hemorrhage and a vascular phenotype of *dyrk1aa*^*krb1*^ mutant embryos

Recently, we reported the generation of the *dyrk1aa*^*krb1*^ mutant that displayed microcephaly and autistic behavioral phenotypes in adults, whereas no distinct morphological defects were observed during embryogenesis (Kim et al, 2017). Detailed inspection of *dyrk1aa*^*krb1*^ homozygous mutant embryos, however, revealed a cerebral hemorrhage phenotype as early as 52 hours post-fertilization (hpf) (arrows in Fig. 1A); 28.1% of the offspring of *dyrk1aa*^*krb1*^ mutants in a clutch displayed cerebral hemorrhage whereas less than 11.7% of wild-type (WT) offspring showed spontaneous hemorrhage at 52 hpf (Fig. 1B). Cerebral hemorrhage was detected as patches in the forebrain, midbrain, hindbrain, and retina, with some embryos exhibiting multiple hemorrhages simultaneously (Fig. 1A). To obtain more pronounced hemorrhagic effects and observe a more explicit phenotype, we challenged the mutant embryos with heat stress by incubation at 35°C for 2.5 hours from 48 hpf to 50.5 hpf and kept them at 28.5°C until 52 hpf. The following experiments for hemorrhagic phenotype were performed in the heat stress condition. It has been previously reported that cardiovascular stress aggravates the hemorrhagic phenotype in the brain, presumably by increasing the heart rate (Barrionuevo & Burggren, 1999; Wang et al, 2014). Consistently, the cerebral hemorrhage of WT and *dyrk1aa*^*krb1*^ mutants under the heat-stressed condition increased by 16.7% and 40%, respectively (Fig. 1B; Heat-stressed).

**Figure 1.**
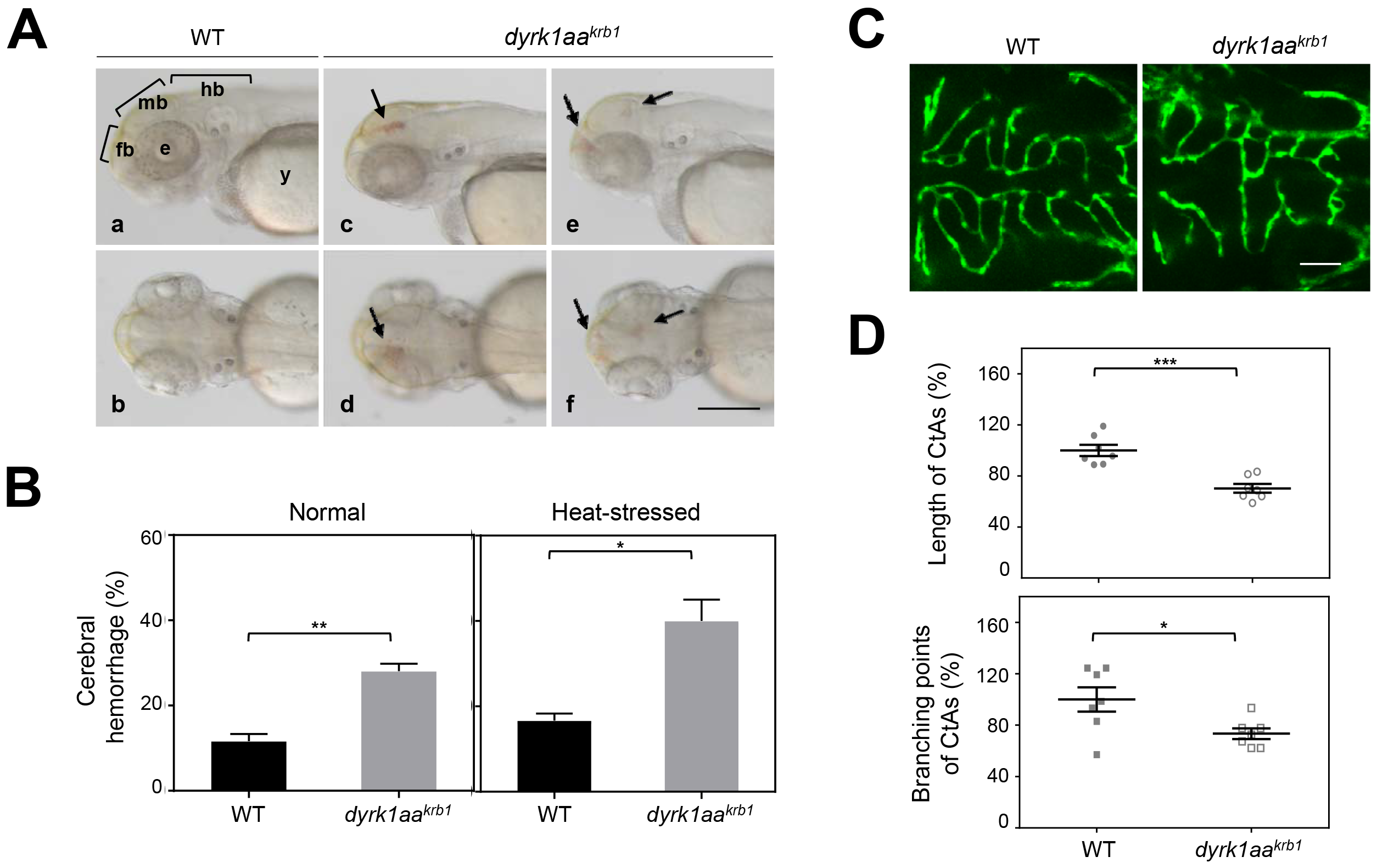
*dyrk1aa*^*krb1*^ mutant embryos show cerebral hemorrhagic phenotype and abnormal development of CtAs (central arteries) in the brain. (**A**) Cerebral hemorrhage was observed in *dyrk1aa*^*krb1*^ mutant embryos (*dyrk1aa*^*krb1*^) at 52 hpf (c-f, arrows), compared to wild type (WT) (a, b). (a, c, e), lateral view; (b, d, f), dorsal view; fb, forebrain; mb, midbrain; hb, hindbrain; e, eye; y, yolk. Scale bar = 250 μm (**B**) Embryonic cerebral hemorrhage of wild type occurred spontaneously within 11.7%, whereas *dyrk1aa*^*krb1*^ embryos showed cerebral hemorrhage of 28.1% penetrance (Normal) counted by *o*-dianisidine staining. The cerebral hemorrhage of wild type and *dyrk1aa*^*krb1*^ mutant increased up to 16.7% and 40%, respectively, by inducing heat-stress with 2.5 hours incubation at 35 °C during 48-50.5 hpf (Heat-stressed). The mean percentage for each genotype was presented from three independent experiments with approximately 20 embryos for each repeat. (**C**) The confocal fluorescent images at 52 hpf showed the development of the CtAs in WT and *dyrk1aa*^*krb1*^ mutant embryos in the *Tg(kdrl:EGFP)* background. Scale bar = 50 μm (**D**) The lengths and branching points of CtAs in *dyrk1aa*^*krb1*^ mutants were reduced down to 70.2% and 73.3%, respectively, compared to the wild type embryos as 100% at 52 hpf. Allp-values from Student’s t-test: *, *p* < 0.05, **, *p* < 0.01, ***,*p* < 0.005; Error bars are ± S.E.M.

Because cerebral hemorrhage is sometimes accompanied by defective cerebrovascular formation (Arnold et al, 2014), we examined the formation of central arteries (CtAs) in the hindbrain, a well-characterized, stereotypical developing cerebrovascular structure using *Tg(kdrl:EGFP)* transgenic animals (Ulrich et al, 2011) in the WT or *dyrk1aa*^*krb1*^ mutants. CtAs sprouted from the primordial hindbrain channels, which contained the pool of endothelial cells required for CtA formation, and invaded the hindbrain between 32 and 36 hpf with a stereotypical morphology within rhombomeres formed with over 50% ipsilateral CtA connectivity at 48 hpf (Bussmann et al, 2011; Ulrich et al, 2011). A detailed, high-resolution confocal imaging analysis to compare the CtA formation of *dyrk1aa*^*krb1*^ mutants and WT at 52 hpf revealed that the stereotypical structure of CtAs was abolished in *dyrk1aa*^*krb1*^ mutant embryos (Fig. 1C). To quantitate the CtA vascular defects, the length and branching points of CtAs were measured, representing the migration/proliferation and sprouting activities of CtA endothelial cells, respectively (AlMalki et al, 2014; Phng et al, 2009; Timar et al, 2001)(Fig. 1D). In *dyrk1aa*^*krb1*^ mutants, the total lengths and branching points of CtAs were reduced to 70.2% and 73.3% compared to the WT control, respectively (Fig. 1D). In the following experiments, we used the length and branching points of CtAs as quantitative measures of defects in vascular development.

CtA angiogenic defects were also confirmed by examining RNA expression of vascular markers *kdr1 (vegfr2)* and *dll4* by whole-mount RNA *in situ* hybridization (WISH) at various developmental stages including 30, 36 hpf, and 52 hpf (Fig. S1; *kdrl* and *dll4*). Consistent with the angiogenic defects revealed by *Tg(kdrl:EGFP)* transgenic animals, expression of *kdrl* and *dll4* in the vasculature of the hindbrain was reduced in *dyrk1aa*^*krb1*^ mutant embryos at all stages examined (red arrows in Fig. S1; *kdrl* and *dll4*). These vascular defects in mutants appeared not to be due to gross defects of brain development because the expression of *krox20*, which marks rhombomere boundaries (Moens & Prince, 2002; Oxtoby & Jowett, 1993), and *isl1*, which labels primary motoneurons in the hindbrain (Ericson et al, 1992; Inoue et al, 1994; Korzh et al, 1993), were grossly unaffected in mutant embryos (Fig. S1; *krox20* and *isl1*). These vascular defects did not appear to be due to defective heart development or reduced blood flow, based on the normal heart pumping rate of mutants compared to the WT embryos (Fig. S2).

### The *dyrk1aa* gene is expressed in endothelial cells and the nervous system during development

We examined transcripts of *dyrk1aa* in zebrafish embryos by WISH during embryogenesis. The *dyrk1aa* mutant had a broad expression in the forebrain (black arrowheads/brackets), midbrain (gray arrowheads/brackets), hindbrain (blue arrowheads/brackets), spinal cord (orange arrowheads) heart (asterisks), and retina (red arrows) at 24, 48, and 72 hpf (Fig. 2Aa-l). Transverse sections of WISH embryos at the midbrain level showed that *dyrk1aa* was broadly expressed in the tectum (green arrows), tegmentum (black arrow), the ganglionic cell layer (purple arrows), and the inner nuclear layer of the retina (red arrows) (Fig. 2A; m and n). We also examined the temporal expression patterns of *dyrk1aa* mRNA according to developmental stages by reverse transcription-PCR (RT-PCR). Transcripts of *dyrk1aa* mRNA were strongly detectable at the one-cell stage but decreased after 6 hpf due to the maternal effect (Harvey et al, 2013; Mathavan et al, 2005) (Fig. 2B). Zygotic *dyrk1aa* expression appeared to start after 15 hpf, increasing at 24 hpf, and strong expression was maintained until 5 days post-fertilization (dpf) (Fig. 2B).

**Figure 2.**
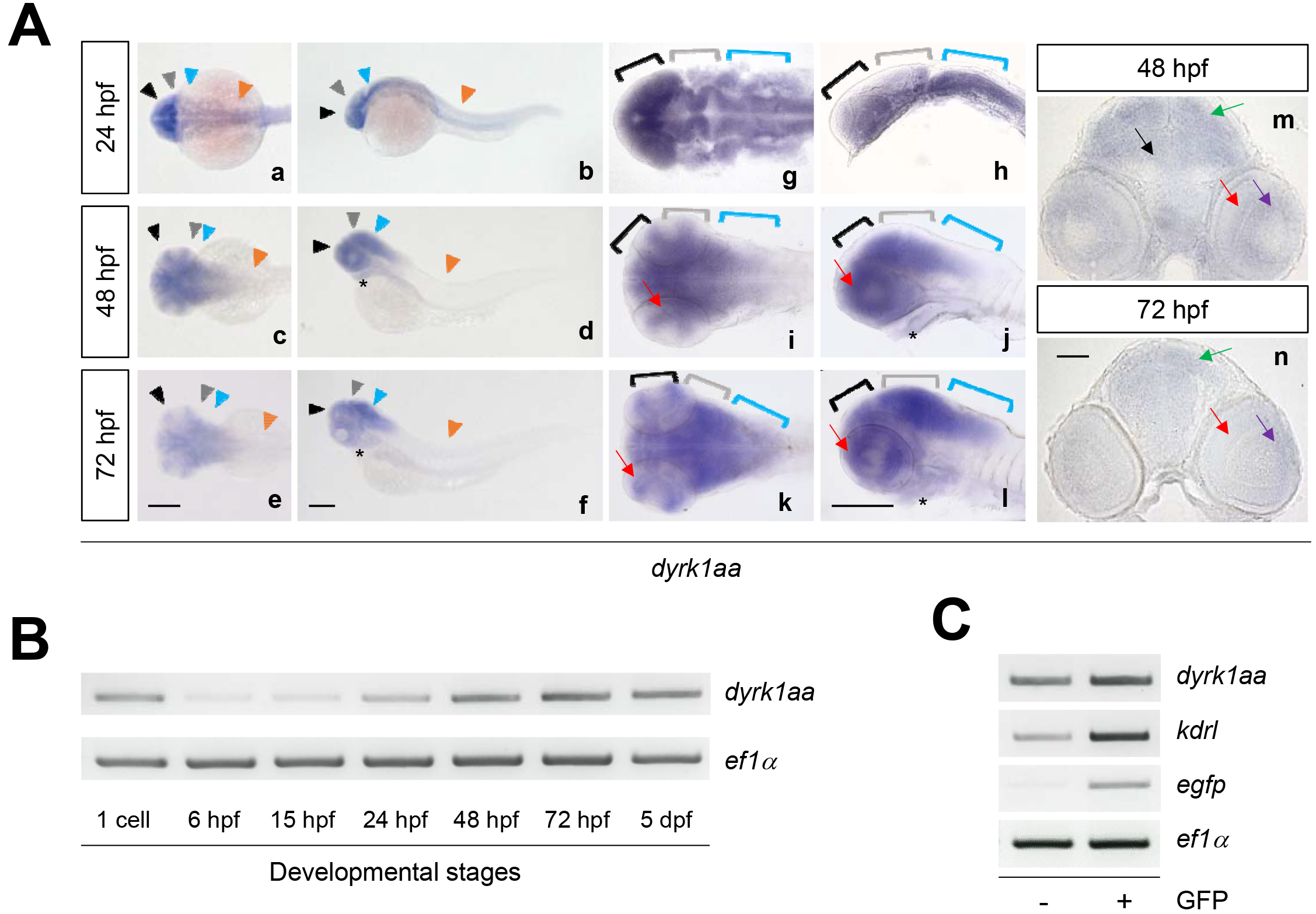
Zebrafish *dyrk1aa* is expressed in the developing brain during embryogenesis and detected in endothelial cells of the vasculature. (**A**) WISH (whole mount *in situ* hybridization) showed that *dyrk1aa* was expressed in the forebrain (black arrowheads, a-f; black brackets, g-l), the midbrain (gray arrowheads, a-f; gray brackets, g-l), the hindbrain (blue arrowheads, a-f; blue brackets, g-l), and the spinal cord (orange arrowheads, a-f) at 24, 48 and 72 hpf. It was also detected in the heart (asterisks, d, f, j and l) and in the retina (red arrows, i-l) at 48 and 72 hpf. (m and n) Sectioned images of WISH embryos showed the expression of *dyrk1aa* in the tectum (green arrows), tegmentum (black arrow), the ganglionic cell layer (purple arrows), and the inner nuclear layer of the retina (red arrows) at 48 hpf and 72 hpf. Scale bars = 200 μm in (a-l) and 50 μm in (m, n) (**B**) Semi-quantitative RT-PCR analysis of *dyrk1aa* mRNA expression in whole embryos at developmental stages. *dyrk1aa* expression was observed at the one cell stage followed by diminishing at 6 hpf and resumed after 24 hpf. (**C**) Semi-quantitative RT-PCR analysis using endothelial cells isolated by FACS for sorting GFP-positive and -negative cells of *Tg(kdrl:EGFP)* embryos at 48 hpf revealed that *dyrk1aa* mRNA was expressed in GFP-positive endothelial cells as well as GFP-negative embryonic cells.

We also examined whether *dyrk1aa* mRNA was specifically expressed in endothelial cells by performing RT-PCR analyses using GFP-positive *Tg(kdrl:EGFP)* endothelial cells isolated by fluorescence-activated cell sorting. The *dyrk1aa* mRNA was enriched in GFP-positive endothelial cells but was also expressed in GFP-negative cells, suggesting a role in endothelial as well as non-endothelial cells (Fig. 2C). These expression patterns were consistent with the reported mammalian *Dyrk1A* expression in the heart primordium and the central nervous system of developing mouse embryos, including the inner (neural) layer of the optic cup(Hammerle et al, 2008; Rahmani et al, 1998), and endothelial cells isolated from *Dscr1* transgenic mice, as assessed by western blotting (Baek et al, 2009), suggesting a functional conservation across species.

### Rescuing *dyrk1aa*^*krb1*^ mutant phenotypes by *dyrk1aa* expression

To confirm that loss of function of the *dyrk1aa* gene was responsible for *dyrk1aa*^*krb1*^ mutant phenotypes, we tested whether WT *dyrk1aa* mRNA rescued the cerebral hemorrhage and aberrant vascular phenotype in *dyrk1aa*^*krb1*^ mutants by globally expressing full-length WT *dyrk1aa* mRNA. Using o-dianisidine staining that detects hemoglobin activity and allows observation of cerebral hemorrhage more closely, the high incidence of cerebral hemorrhage in *dyrk1aa*^*krb1*^ mutants under heat stress (41.7% of the offspring) was shown to be significantly reduced down to 25.4% by 0.1 ng *dyrk1aa* mRNA injection (Fig. 3A, 3B), whereas the same dose injected into the WT background had little effect (Fig. 3A, 3C). Similarly, the reduced mean percentages of lengths (66.5%) and branching points (61.2%) of CtAs in *dyrk1aa*^*krb1*^ mutants relative to WT controls (100%) were rescued (89.9% and 122%, respectively) with the expression of 0.1 ng *dyrk1aa* mRNA (Fig. 3D, 3E).

**Figure 3.**
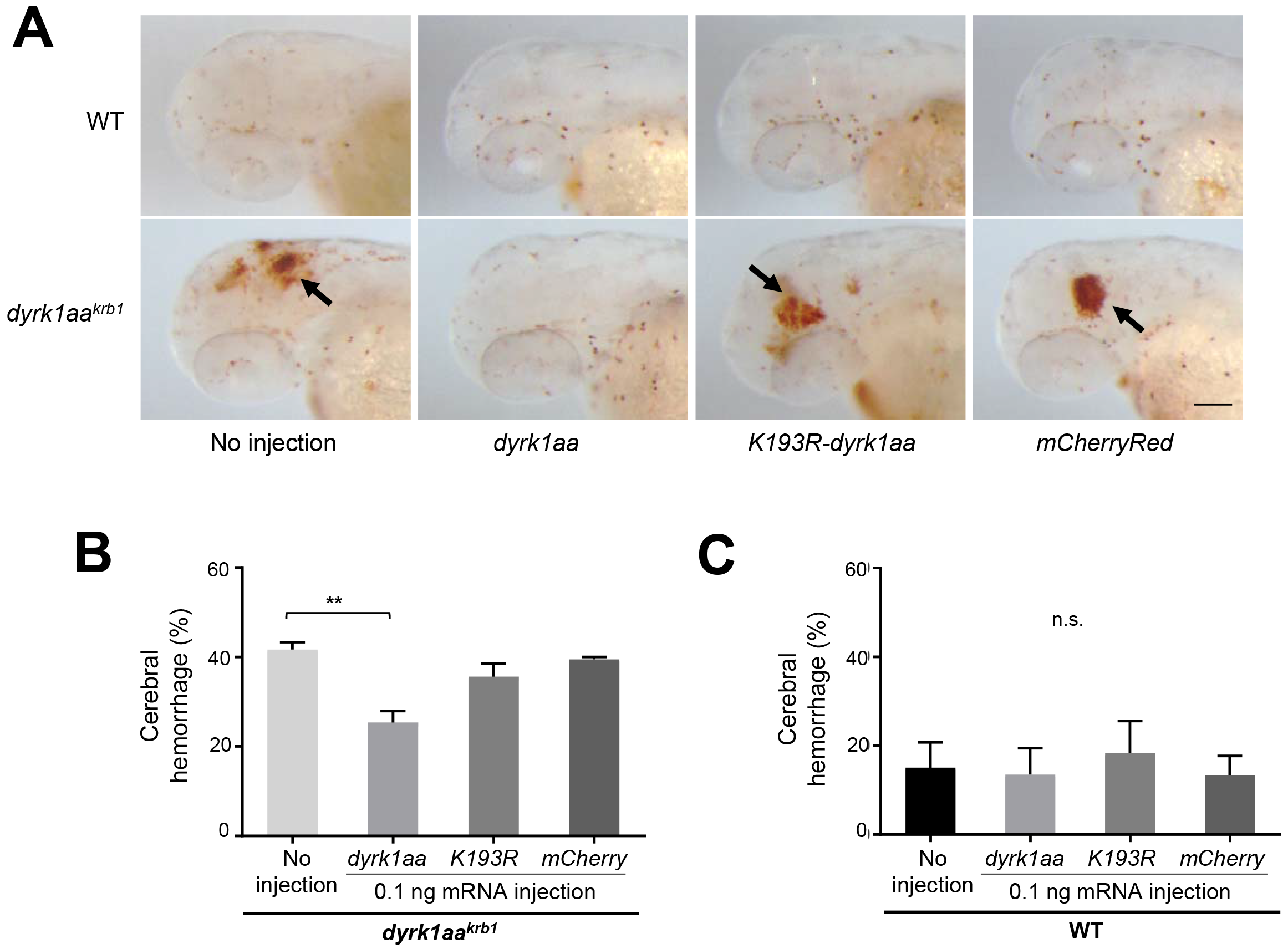
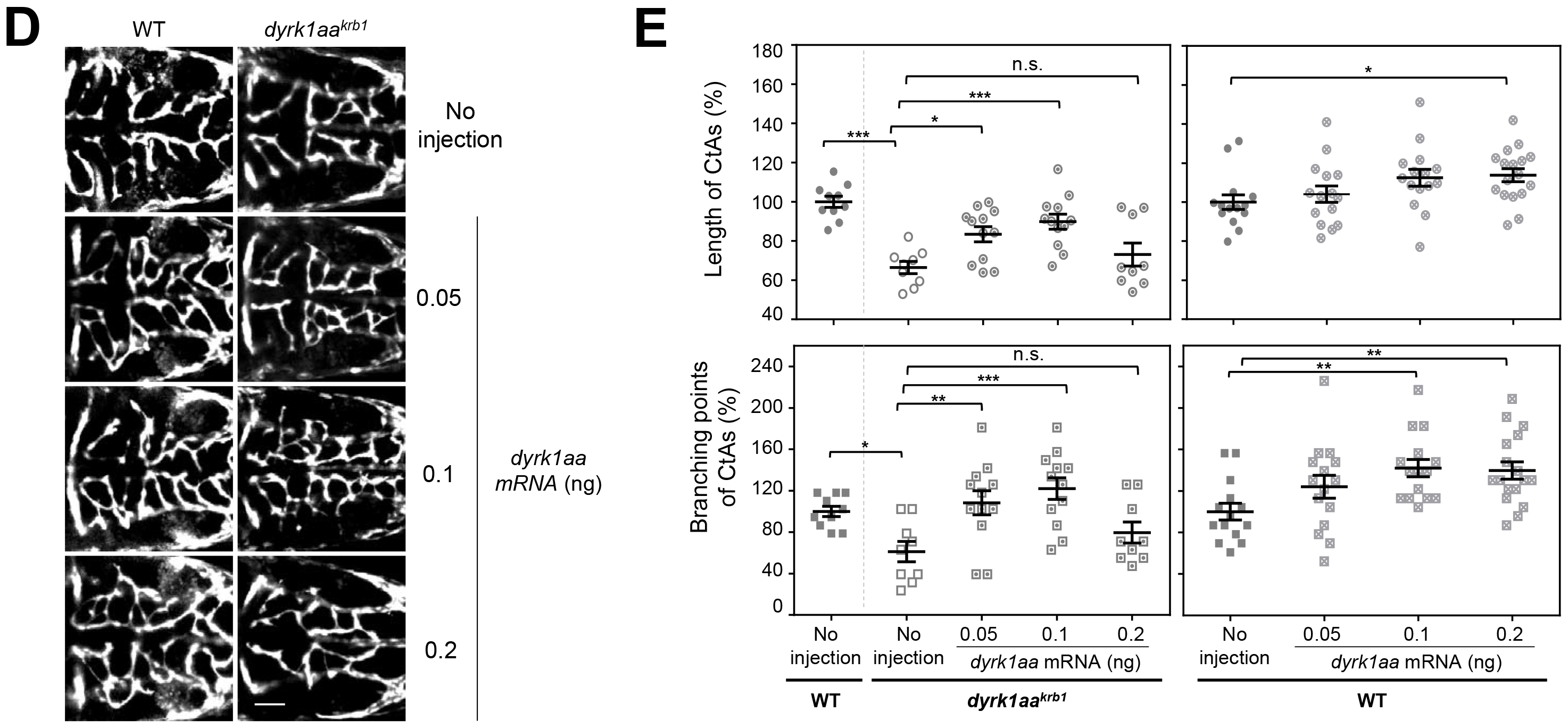
The cerebral hemorrhage and CtA angiogenic defects in *dyrk1aa*^*krb1*^ mutant can be rescued by *dyrk1aa*, with the kinase activity of Dyrk1aa are required for its phenotypic rescue. (**A**) The *o*-dianisidine staining images at 52 hpf showed that the cerebral hemorrhage (arrows) of *dyrk1aa*^*krb1*^ embryos was rescued by wild type *dyrk1aa* mRNA injection, but not in noinjection control (no injection), *Kl93R-dyrk1aa*, or *mCherryRed*mRNA. Scale bar = 100 μm. (**B**) The cerebral hemorrhage of *dyrk1aa*^*krb1*^ was reduced from 41.7% to 25.4% by injecting wild type *dyrk1aa* mRNA of 0.1 ng, but not significantly changed by *K193R-dyrk1aa* and *mCherryRed* mRNA at 52 hpf. (**C**) Injection of mRNAs of wild type *dyrk1aa, K193R-dyrk1aa* or *mCherryRed* control did not affect the cerebral hemorrhage in WT embryos at 52 hpf. The mean percentage for each genotype was presented from three independent experiments with approximately 20 embryos in each repeat. (**D**) The compiled images of CtAs of *Tg(kdrl:EGFP)* at 52 hpf by confocal microscopy showed that the angiogenic defects of the CtAs of *dyrk1aa^krb1^* embryos was also rescued by WT *dyrk1aa* mRNA injection in a dose-dependent manner. Scale bar = 50 μm (**E**) The mean percentages of length and branching points of CtAs in *dyrk1aa^krb1^* mutants are rescued from 66.5% to 83.4% and from 61.2% to 108.3%, respectively, with 0.05 ng of *dyrk1aa* mRNA injection, relative to WT as 100 %, while rescued to 89.9% and 122.0%, respectively, with 0.1 ng of *dyrk1aa* mRNA injection. Expression with 0.2 ng of *dyrk1aa* mRNA did not exhibit the rescue effect. Expression of *dyrk1aa* mRNA with same doses in WT embryos also increased the length and branching points of CtAs (113.8% and 139.6% with 0.2 ng of *dyrk1aa* mRNA, respectively). *p*-values by one way ANOVA: \**p* < 0.05, \*\**p* < 0.01,\*\*\**p* < 0.005; n.s., not significant. Error bars are ± S.E.M.

Dyrk1a regulates its target substrate proteins via phosphorylation (Galceran et al, 2003; Hammerle et al, 2003). To verify whether the cerebral hemorrhage and CtA defects in *dyrk1aa*^*krb1*^ mutants were dependent on the kinase activity of Dyrk1aa, we performed rescue experiments with the *K193R-dyrk1aa* mRNA, the predicted kinase-dead form of Dyrk1aa (Himpel et al, 2001). *K193R-dyrk1aa* mRNA expression failed to rescue the defects of hemorrhage and CtA formation with comparable doses of WT-*dyrk1aa* mRNA (Fig. 3B, Fig. S3A). These results suggest that the kinase activity of Dyrk1aa is critical for normal CtA development and prevention of hemorrhage.

### Ultrastructural analyses of cerebral vessels in *dyrk1aa*^*krb1*^ mutants by transmission electron microscopy

To examine whether the *dyrk1aa* mutation caused an ultrastructural change in brain vessels, we analyzed the cytoarchitecture of blood vessels in the WT and *dyrk1aa*^*krb1*^ embryos at 52 hpf using transmission electron microscopy. In the WT group, blood vessels composed of endothelial cells and lumens were well formed and tightly arranged at this stage (Fig. 4A and A’), although smooth muscle cells, pericytes, and astrocytes were in the process of differentiation and not yet clearly identified (Liu et al, 2007). Characteristically, brain tissues in *dyrk1aa*^*krb1*^ mutants exhibited enlarged interstitial spaces presumably due to loose connections between layers of vessel walls (arrowheads in Fig. 4B and B’), suggesting that this abnormal formation of vessel walls was one of the causes of the hemorrhagic phenotype in *dyrk1aa*^*krb1*^ mutant embryos.

**Figure 4.**
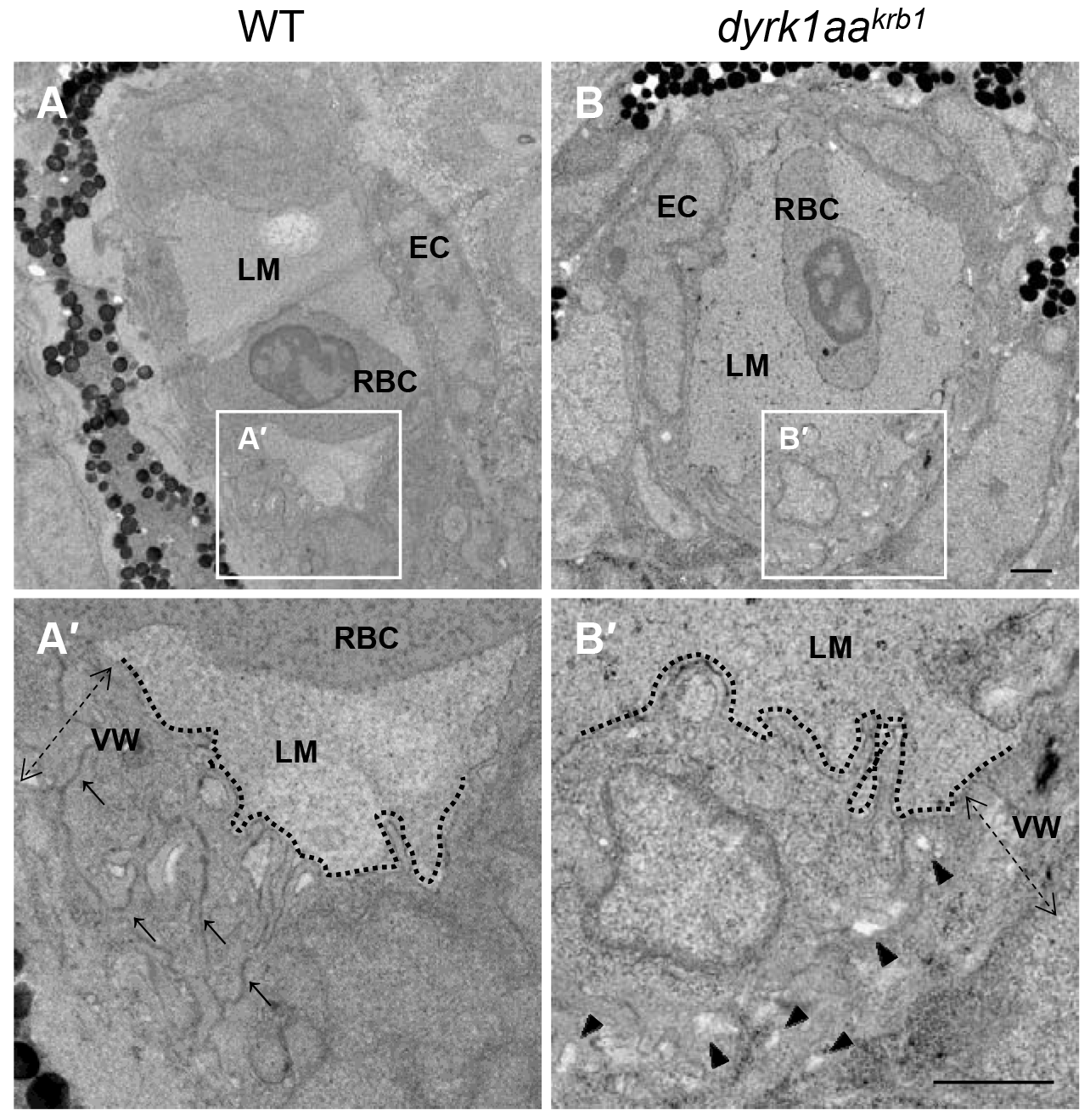
The transmission electron microscopy reveals that *dyrk1aa*^*krb1*^ embryos have abnormal vessel walls in the brain at 52 hpf. (**A, B**) The brain vessels in WT and *dyrk1aa*^rb1^. A’ and B’ indicate the enlarged images of white boxes in A and B. (**A’, B’**) Arrows in A’ indicate the compact structure of vessel walls, whereas arrowheads in B’ designate the loose connection of vessel walls. Dashed lines demarcate the border between the vessel walls and the lumen. EC, endothelial cell; LM, vessel lumen; RBC, red blood cell; VW (doted arrows), vessel wall. Scale bars =1 μm.

### Dyrk1a inhibition by harmine and chemical screening

Cerebral angiogenic defects and hemorrhage in *dyrk1aa*^*krb1*^ mutants were also recapitulated by harmine (7-methoxy-1-methyl-9H-pyrido[3,4-b]-indole), a well-known DYRK1A inhibitor(Bain et al, 2007; Gockler et al, 2009). WT zebrafish embryos exposed to different concentrations of harmine (10, 25, and 50 μM) starting at 24 hpf for 28 hours showed the hemorrhage and CtA formation defects at 52 hpf, quite similar to those of the *dyrk1aa*^*krb1*^ mutants, and the number of CtA sprouts was reduced down to 15.2% of the control number at a concentration of 50 μM harmine (Fig. 5A, B). The harmine-induced hemorrhagic phenotype increased up to 65.7% that of the offspring, compared to 2.3% of the DMSO-treated controls (Fig. 5C). This chemical suppression of Dyrk1aa using harmine further suggested that the vascular phenotypes of *dyrk1aa*^*krb1*^ mutants were due to the loss of Dyrk1aa function.

**Figure 5.**
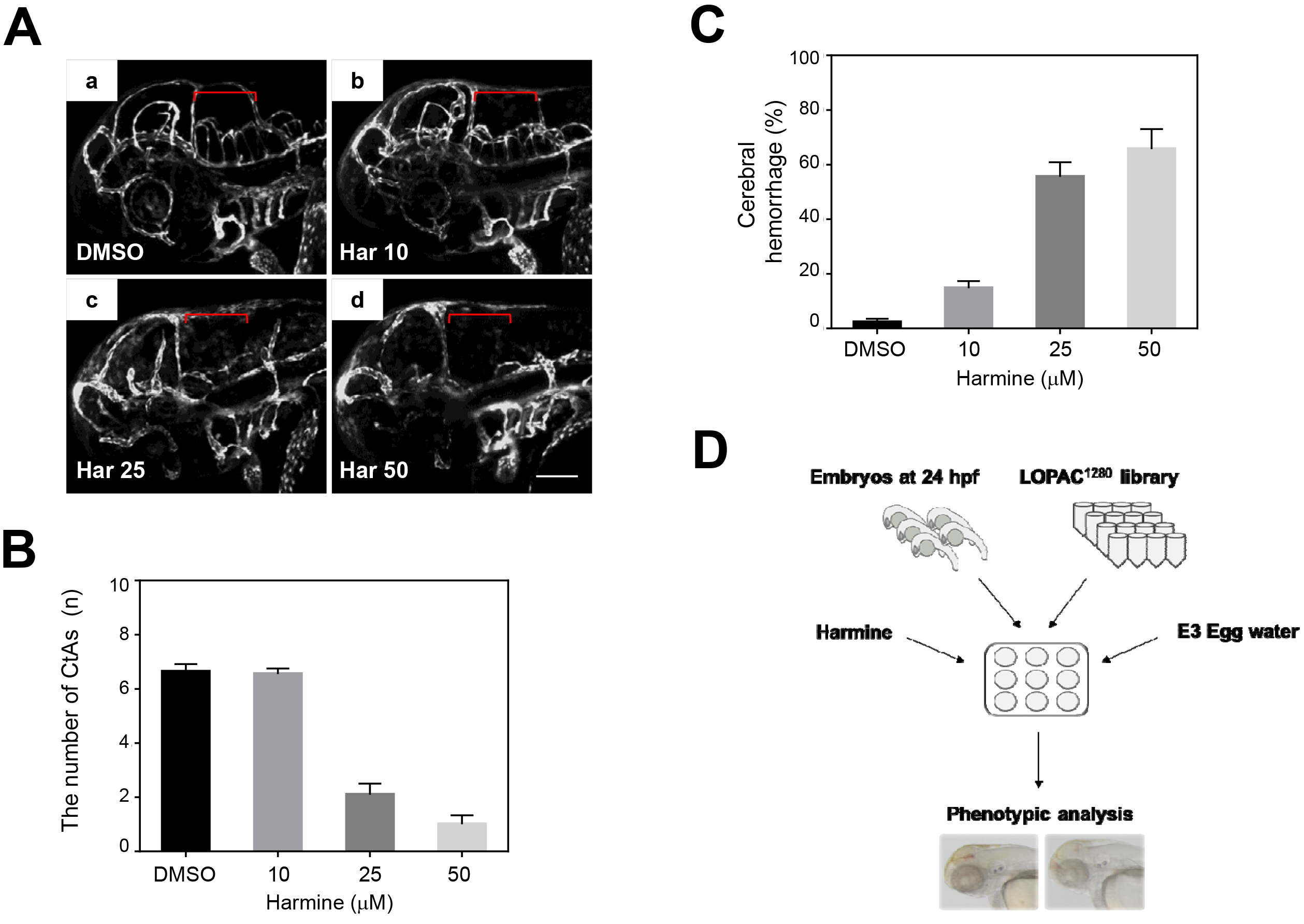
Inhibition of Dyrk1aa by harmine induces the brain hemorrhage and the vasculature defects in the hindbrain. (**A**) WT *Tg(kdrl:EGFP)* embryos were treated with increasing concentrations of harmine, a DYRK1A inhibitor: a, DMSO; b, c, d, 10 μM; 25 μM; 50 μM of harmine, respectively, from 24 hpf until 52 hpf. Vascular patterning defects of CtAs were shown (red brackets) by confocal imaging at 52 hpf (lateral view). Scale bar = 100 μm (**B**) Quantification of the number of developing CtAs by harmine treatment. The numbers of CtAs were dramatically reduced by treating harmine in a dose-dependent manner. *N* = 11 each group. (**C**) Quantification of the brain hemorrhage penetrance. The cerebral hemorrhagic phenotype of 2.3% in DMSO-treated embryos was increased from 14.8% up to 65.7% by treating harmine from 10 μM to 50 μM. The mean percentage for each treatment was presented from three independent experiments with approximately 40 embryos in each repeat. (**D**) Schematic drawing shows the strategy of the *in vivo* chemical library screening to identify small molecule modifiers for cerebral hemorrhagic phenotype upon DYRK1A inhibition.

Based on the hemorrhagic phenotype induced by harmine treatment, we developed an embryonic screening assay using a chemical library that consisted of 1,280 FDA-approved and pharmacologically active compounds (LOPAC 1280; Sigma-Aldrich, St. Louis, MO, USA), which allowed us to identify small molecules that modulated the harmine-induced hemorrhagic phenotype. Five WT zebrafish embryos at 24 hpf were placed in each well of a 48-well plate, exposed to 30 μM harmine together with the individual chemicals of the chemical library at 10 μM as a final concentration for 28 hours, and analyzed for the increased or decreased hemorrhagic phenotype (Fig. 5D). As a result, 171 of 1,280 compounds tested were found as phenotype modifiers, which were categorized according to their common features as “class” based on their known functions (Table 1). Some chemicals have already been reported to cause hemorrhage. For example, atorvastatin, which increased the hemorrhage in our screening, has been previously reported to induce hemorrhagic stroke as a side effect in zebrafish embryos (Gjini et al, 2011). Of the 171 compounds, EGTA, a specific calcium chelator, was identified as one of the most efficient suppressors of the hemorrhagic phenotype induced by harmine treatment, in our chemical screening.

**Table 1.**
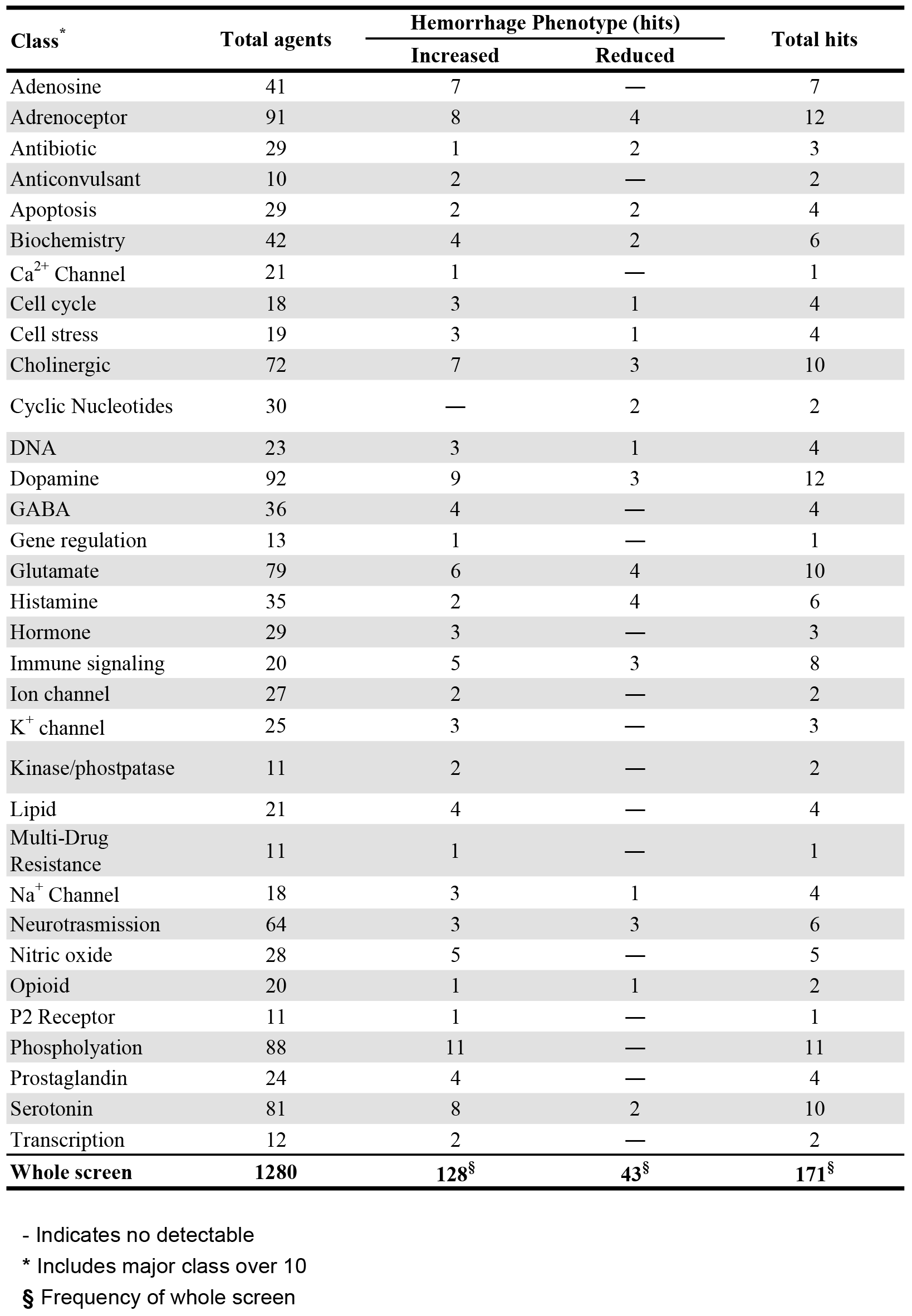
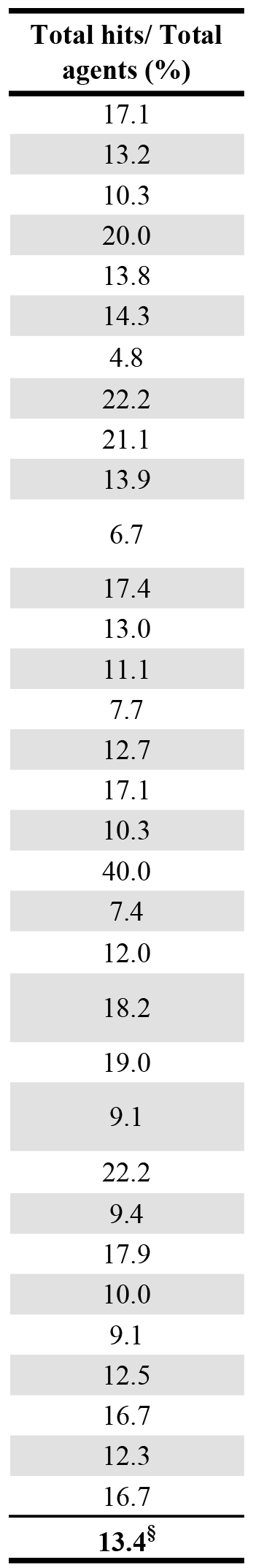
The summary of the in vivo chemical library screening using zebrafish embryos to identify small molecule hits capable of modifying brain hemorrhage phenotype. The table shows high-ranked pharmacologic classes of 171 molecules that modify the hemorrhage phenotype induced by harmine treatment.

### EGTA effectively suppressed the vascular defects of *dyrk1aa*^*krb1*^ mutants

To determine if our findings from the chemical screening were applicable to the genetic model, we added EGTA to the *dyrk1aa*^*krb1*^ mutants and examined the hemorrhagic and CtA development. Under heat-stressed conditions, the EGTA treatment significantly reduced the hemorrhagic phenotype of *dyrk1aa*^*krb1*^ mutants at a specific concentration of 10 nM (from 37.5% to 23.6%; Fig. 6A, 6B; *p* < 0.05). In addition, a reduced CtA development of mutants in lengths and branching points (76% and 57.4% reduction compared to the wild type, respectively) was also rescued up to 86.6% and 85.9% of the normal levels, respectively, by treatment with the same concentrations of EGTA (Fig. 6C, 6D; *p* < 0.01). EGTA treatment appeared to be effective only within a narrow range, because 1 nM or 100 nM EGTA treatment failed to rescue the vascular defects of *dyrk1aa*^*krb1*^ mutants, except for the rescue of CtA branching points with 1nM EGTA (Fig. 6B, 6D). In contrast, EGTA treatment of the WT embryos did not induce any vascular defects (Fig. S4), suggesting a specific role of EGTA on *dyrk1aa*^*krb1*^ mutants.

**Figure 6.**
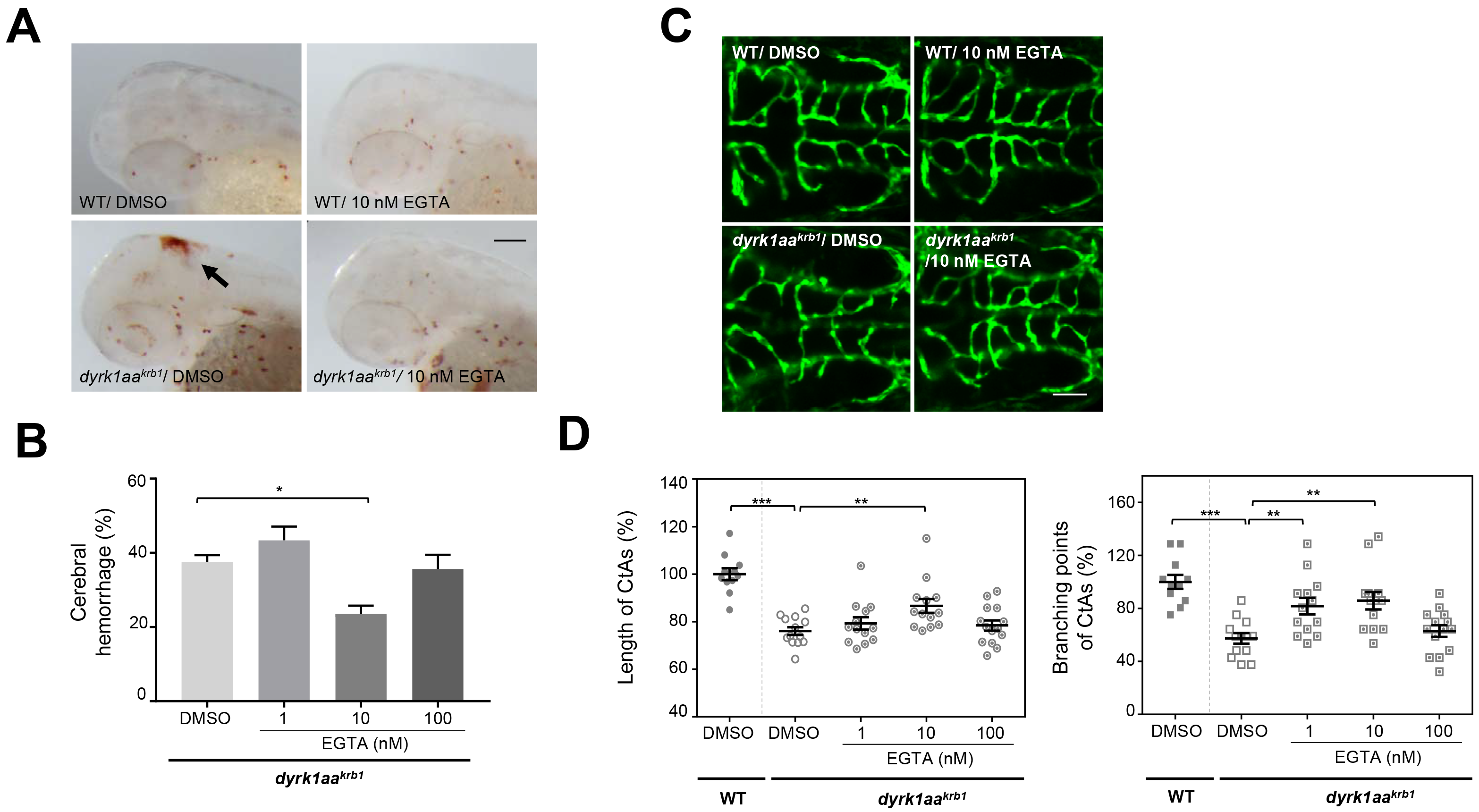
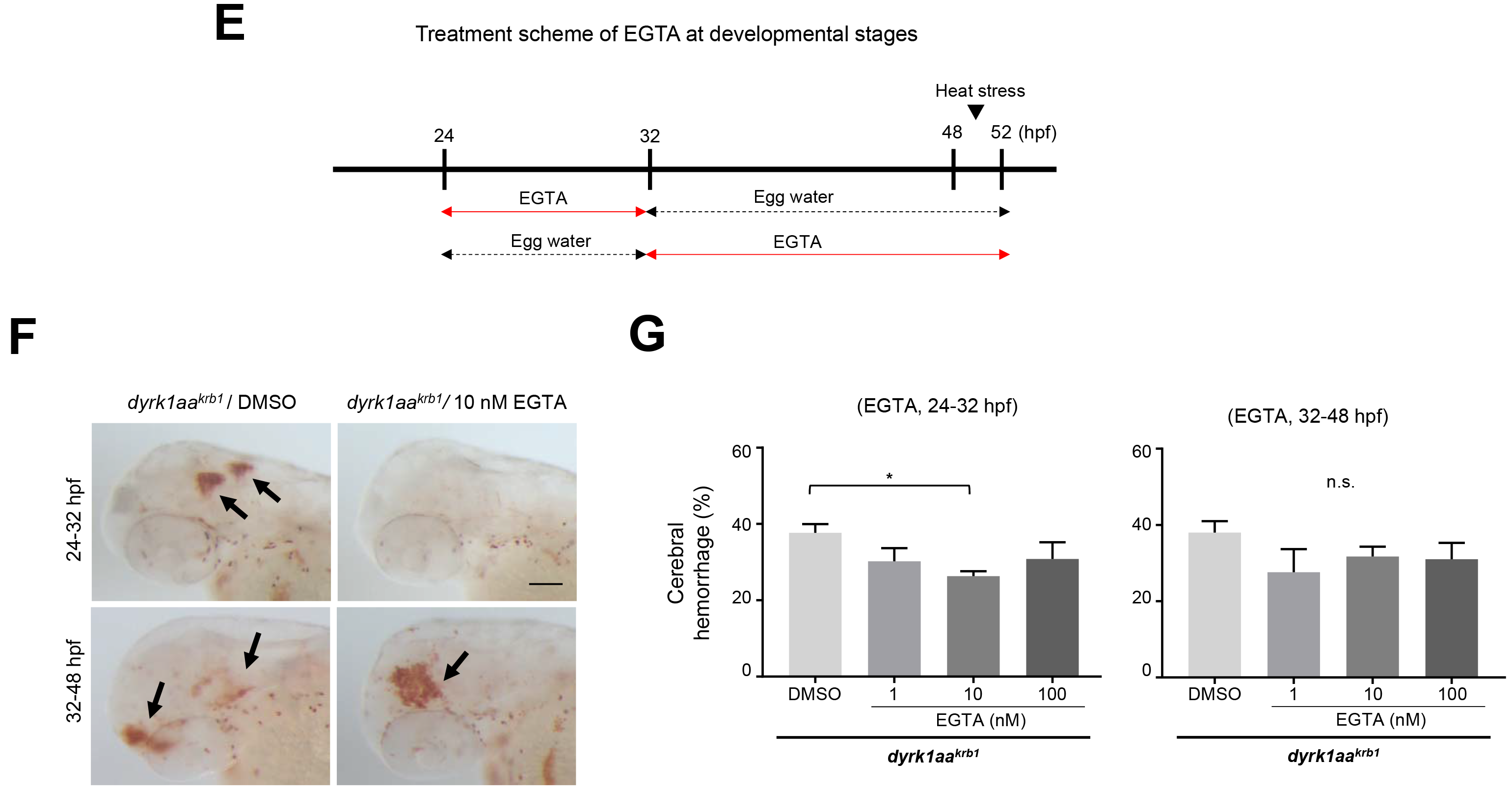
EGTA identified by the chemical library screening effectively rescues the cerebral hemorrhage and abnormal CtA development of *dyrk1aa*^*krb1*^ mutants. (**A**) The o-dianisidine staining images at 52 hpf showed that the cerebral hemorrhage (arrow) of *dyrk1aa*^*krb1*^ embryos are rescued by treating 10 nM EGTA. WT embryos were not affected by the treatment of the same concentration of EGTA. (**B**) Quantitation of the rescue of the cerebral hemorrhagic phenotype by EGTA. The cerebral hemorrhage in *dyrk1aa*^*krb1*^ embryos are reduced from 37.5% to 23.6% by 10 nM EGTA treatment. The data is presented with five independent experiments with approximately 20 embryos in each repeat. (**C**) Compiled confocal microscopy images of the rescue effect on angiogenic defects of CtAs in *dyrk1aa*^*krb1*^ embryos by EGTA treatment. (**D**) Quantitative data showed that the reduced mean percentages of length and branching points of CtAs in *dyrk1aa*^*krb1*^ embryos at 52 hpf (76 % and 57 %, respectively compared to WT embryos as 100 %) were increased up to 86.6 % and 85.9 %, respectively by 10 nM EGTA treatment. *N* ≥ 11 each group. (**E**) The bar diagram depicts the treatment scheme of EGTA at developmental stages from 24 to 52 hpf in *dyrk1aa*^*krb1*^ embryos. (**F**) The cerebral hemorrhage of *dyrk1aa*^*krb1*^ embryos at 52 hpf (arrows) were rescued by 10 nM EGTA treatment for 24-32 hpf but not by the treatment for 32-48 hpf. (**G**) Quantitative data showed that the cerebral hemorrhage of *dyrk1aa*^*krb1*^ embryos were reduced from 37.7% to 26.3% by treating 10 nM EGTA for 24-32 hpf. Scale bars in **A, F** = 100 μm; in **C** = 50 μm. *p*-values by one-way ANOVA: *, *p* < 0.05, **, *p* < 0.01, ***, *p* < 0.005; n.s., not significant. Error bars are ± S.E.M.

To identify the temporal requirement of EGTA for the suppressive effects, developing embryos were incubated with 10 nM EGTA during an early period of 8 hours (24~32 hpf), followed by washing, or during a late period of 16 hours (32~48 hpf) (Fig. 6E). Interestingly, only the early treatment with EGTA was effective in significantly preventing the cerebral hemorrhage (from 37.7% to 26.3%), whereas the late treatment was not (Fig. 6F, 6G). Because sprouting and elongation of cerebral vessels for angiogenesis occurs actively during day 1 post-fertilization(Fujita et al, 2011; Isogai et al, 2001), EGTA may exert its suppressive effects on the vascular defects of *dyrk1aa*^*krb1*^ mutants by regulating early angiogenic processes.

### Transcriptomic analyses of *dyrk1aa*^*krb1*^ mutants

Identification of EGTA as a suppressor of vascular defects in *dyrk1aa*^*krb1*^ mutants by chemical screening implies that calcium signaling may be compromised in the *dyrk1aa*^*krb1*^ mutants. To corroborate this finding, we examined transcriptomic changes of *dyrk1aa*^*krb1*^ mutants compared to WT embryos at 48 hpf by RNA-seq analyses (Fig. 7). This analysis identified 222 transcripts as differentially regulated genes (DEG), of which 101 were up-regulated and 121 were down-regulated in the *dyrk1aa*^*krb1*^ mutant (more than 2-fold and less than 0.5-fold, respectively; *p* < 0.05). When DEGs were analyzed for enriched biological gene ontology (GO) categories by the functional annotation tools in DAVID (Database for Annotation, Visualization and Integrated Discovery, https://david.ncifcrf.gov), the calcium ion binding category was the most enriched GO molecular function (MF) (Fig. 7B). This GO category includes genes encoding several calcium-dependent adhesion proteins, protocadherin (*Pcdh*) family members (*Pcdh1g1*, *Pcdh2ab10*, *Pcdh1gc6*, *Pcdh2g17*, *Pcdh1g18*, and *Pcdh1g30*), and calcium-dependent calpains (*capn8 and capn2l*)(Khorchid & Ikura, 2002) (Fig. 7B). Other genes encoding *Myl4* (myosin light chain 4), *Ldlrb* (low density lipoprotein receptor b), and *Masp2* (mannan-binding lectin serine protease 2) in this GO term are also regulated by calcium signaling, directly or indirectly (Kang et al, 1999; Orr et al, 2016; Zhao & Michaely, 2009) (Fig. 7A), although their roles in regulating vascular formation are not understood at present. In addition, the “oxidation-reduction” process, the top-ranked GO biological process (BP), is also well known to affect calcium signaling networks (Gorlach et al, 2015; Lounsbury et al, 2000) (Fig. 7C), and the genes belonging to the “homophilic cell adhesion via plasma membrane adhesion molecules” BP category consist of the six *pcdh* family genes “calcium ion binding” MF category (Fig. 7C). Collectively, alterations of genes in “calcium ion binding” and other GO categories may reflect dysregulation of calcium homeostasis in *dyrk1aa*^*krb1*^ mutants, consistent with the rescue effects by EGTA on vascular defects of mutants.

**Figure 7.**
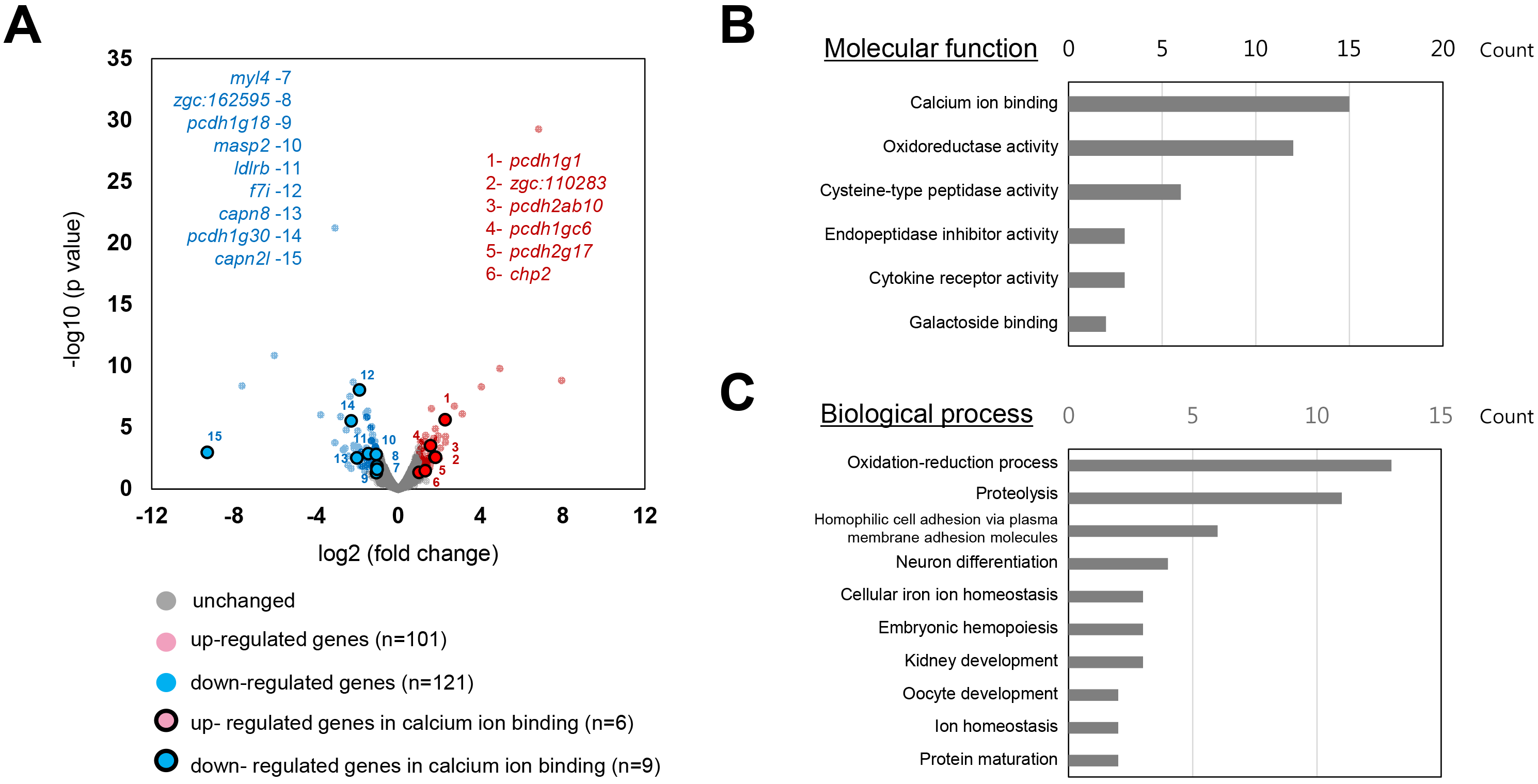
Whole transcriptomic analysis of *dyrk11aa^krb1^* compared with WT by RNA-seq. (**A**) The volcano plot showing the comparison of the whole transcriptomes of the pools of ~20 WT and *dyrk1aa^krb1^* embryos/set at 48 hpf WT and (N=2 biological replication for each group). The colored dots show the differentially up- (red) or down- (blue) regulated genes in *dyrk1aa^Icrb1^* embryos compared to WT embryos (more than 2 fold, *p* < 0.05). The colored dots with black circles indicate the up- (red dot with black circle) and down- (blue dot with black circle) regulated genes in the group of “calcium ion binding” (see text for details). (**B**) A bar graph showing the list of the groups of DEGs (differentially regulated genes) in the classification of the molecular functions. DEGs of the “calcium ion binding” are most abundant. (**C**) A bar graph showing the list of the groups of DEGs in the classification of the biological process. DEGs of the “oxidation-reduction process” are most abundant. *x*-Axis in B and C represents the number of DEG counts in respective annotations.

### Modulation of intracellular calcium signaling also rescued vascular defects of *dyrk1aa*^*krb1*^ mutants

As noted previously, EGTA is a chelating agent that has strong selectivity for calcium ions, primarily by lowering the level of extracellular calcium, eventually affecting the calcium homeostasis inside a cell, which is an essential process for regulating angiogenesis as well as other cellular processes including muscle contraction and neurogenesis (Berridge et al, 2000; Munaron, 2006). To assess whether the modulation of intracellular calcium signaling can recapitulate the effects of EGTA treatment, we attempted to rescue the cerebral hemorrhage and defective CtA formation of *dyrk1aa*^*krb1*^ mutants by inhibiting calcineurin protein with FK506, a well-known specific calcineurin inhibitor(Onuma et al, 1998). The signaling pathway mediated by calcineurin, a serine/threonine protein phosphatase, is one of the major signaling pathways that is regulated by calcium (Crabtree, 2001; Hogan et al, 2003). Interestingly, the hemorrhagic phenotype of *dyrk1aa*^*krb1*^ was rescued by treatment with 50 and 100 ng/ml FK506 in a dose-dependent manner, while no significant change was observed in the WT control when treated (Fig. 8A, 8B). The same concentrations of FK506 also rescued defects in the CtA branching points (100 ng/ml) and length (50 ng/ml) of mutants, although FK506 also increased those of CtAs in the WT control (Fig. 8C, 8D). These data were consistent with the notion that the dysregulation of calcium homeostasis was responsible for the vascular defects of the *dyrk1aa*^*krb1*^ mutants, and such defects could be rescued, at least partially, by manipulating a calcium-dependent signaling pathway.

**Figure 8.**
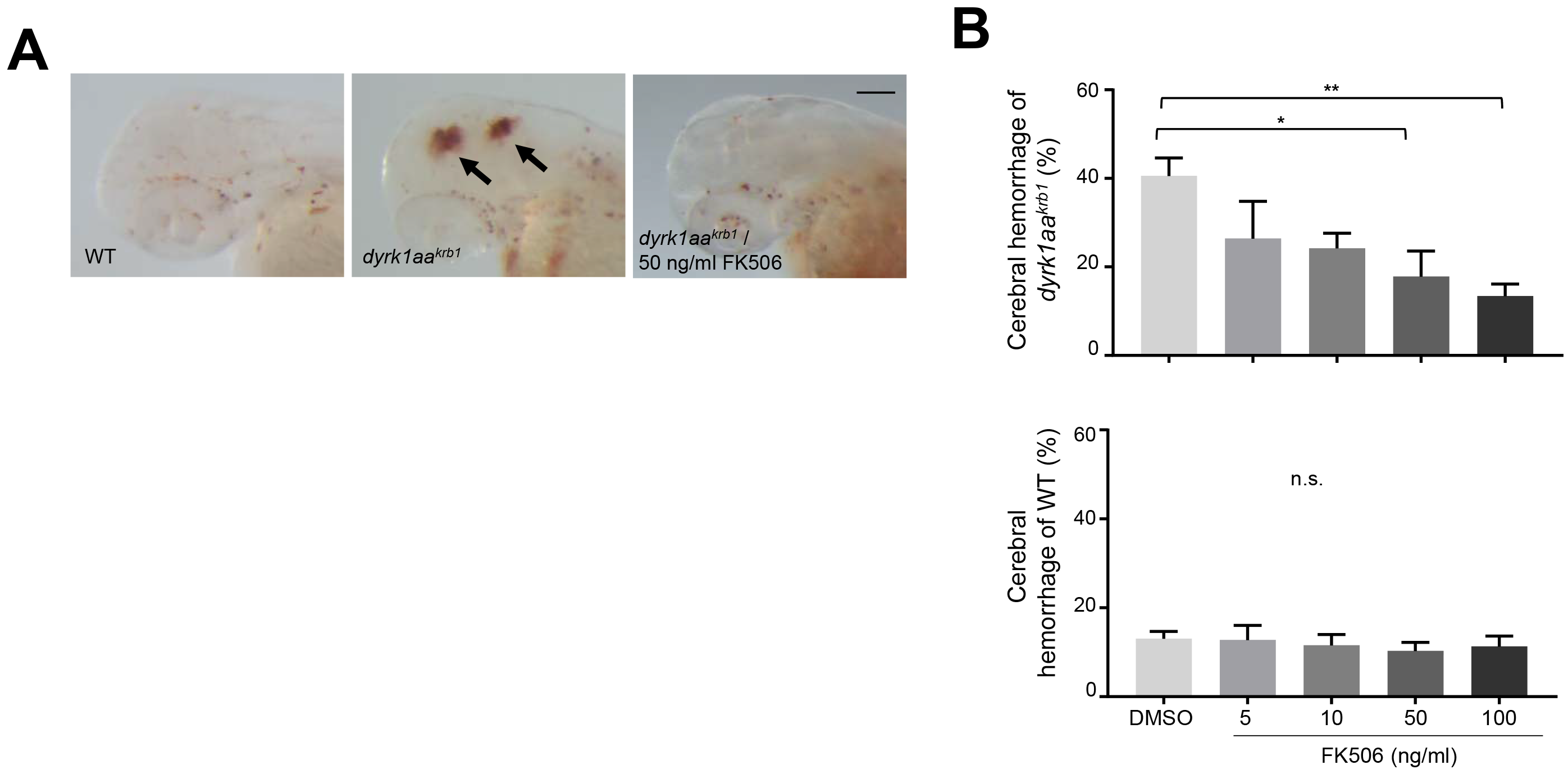
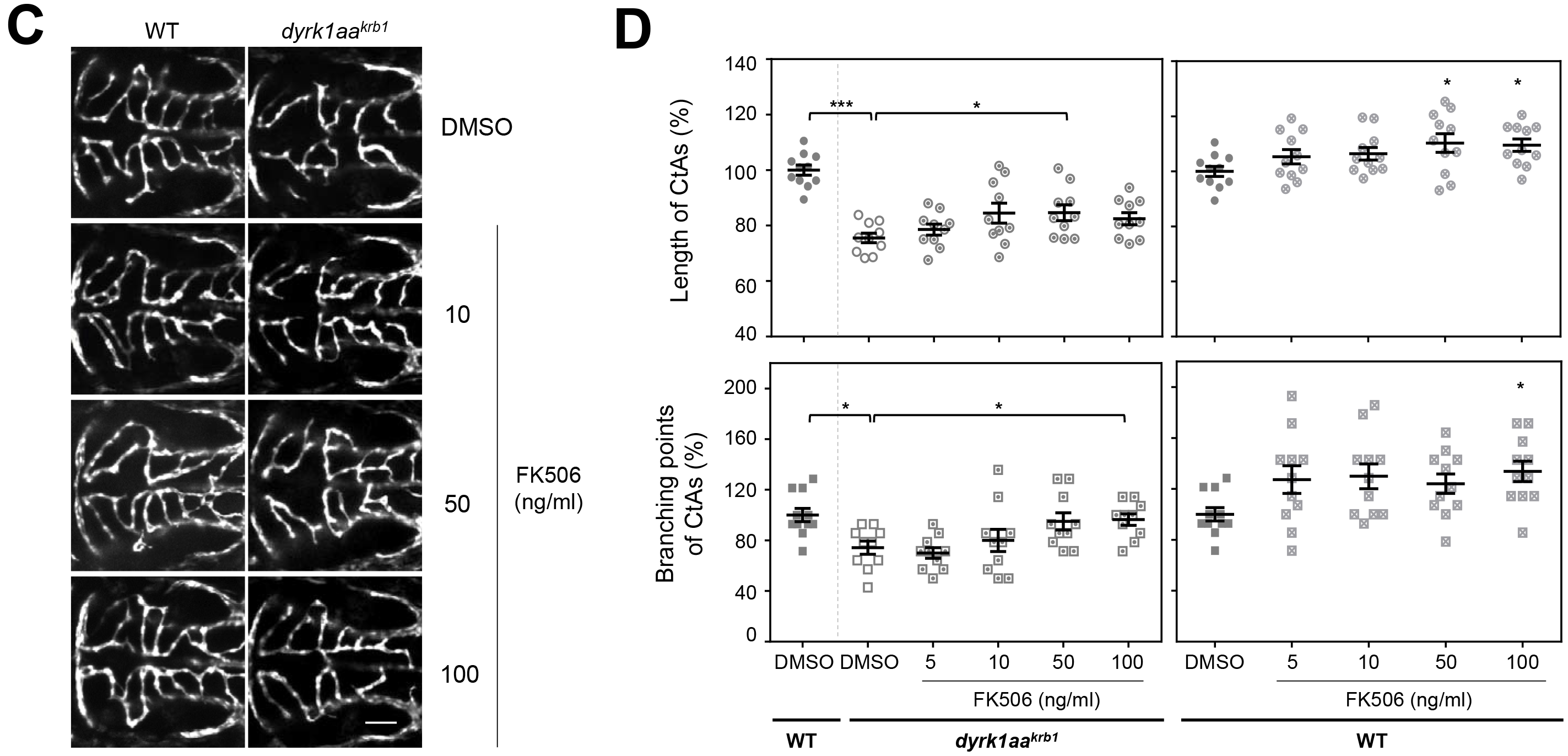
FK506 rescues the cerebral hemorrhage and CtA angiogenic defects in *dyrk1aa*^*krb1*^ mutants. (**A**) The cerebral hemorrhage of *dyrk1aa*^*krb1*^ mutant embryos (arrows) was rescued by the treatment of 50 ng/ml FK506. (**B**) The cerebral hemorrhage in *dyrk1aa*^*krb1*^ mutant embryos at 52 hpf (40.5%) were reduced to 17.8% and 13.4% by the treatment of 50 ng/ml and 100 ng/ml FK506, respectively. No differences was seen in WT by the same treatments. (**C**) The compiled confocal microscopy images of CtAs in the *Tg(kdrl:EGFP)* background showed that CtA defects in *dyrk1aaa^krb1^* embryos were rescued by FK506 treatment in a dose-dependent manner, while CtA angiogenesis in WT were also affected. (**D**) The reduced mean percentage of CtA length of *dyrk1aa*^*krb1*^ mutants (75.5%) was significantly rescued up to 84.6% by 50 ng/ml FK506, while that of mutant branching points (74.3%) to 96.4% by 100 ng/ml FK506 at 52 hpf, compared to WT embryos as 100%. 50 ng/ml FK506 treatment increased the WT CtA length up to 110.3% while 100 ng/ml FK506 treatment increased both length and branching points of WT CtAs (109.5% and 133.8%, respectively). Scale bar in A = 100 μm; in C = 50 μm. *p*-values by one-way ANOVA: *, *p* < 0.05, **, *p* < 0.01, ***, *p* < 0.005; n.s., not significant. Error bars are ± S.E.M.d

## DISCUSSION

Proper cerebrovascular development and maintenance are essential processes for normal brain development and function, and are governed by diverse and coordinated signaling pathways (Hogan & Schulte-Merker, 2017). In this report, we established a less-known function of *DYRK1A*, one of the critical genes responsible for DS, in regulating the angiogenesis and preventing the hemorrhage in the brain, using zebrafish *dyrk1aa* knockout mutants. The *in vivo* chemical screening using zebrafish embryos identified small molecules that were able to modify the cerebral hemorrhage upon Dyrk1A inhibition. Among them, EGTA, a known specific calcium chelator, was identified as one of the most effective small molecules that rescued the vascular defects of *dyrk1aa* mutants. Changes in calcium-related signaling pathways revealed by RNA-seq analyses and the rescuing activity by chemical inhibition of calcineurin, a major calcium-dependent pathway, on the vascular defects corroborated the notion that the vascular defects of *dyrk1aa* mutants were primarily due to calcium dysregulation, which could be reversed by inhibition of excessive calcium-dependent processes.

Zebrafish *dyrk1aa* knockout mutants exhibited compromised vessel integrity with cerebral hemorrhage and angiogenesis simultaneously (Fig. 1, Fig. 4), compared to WT controls. Although the relationship among angiogenesis, vessel permeability, and cerebral hemorrhage during developmental processes is not clearly defined, the interactions of these processes may be important for development of the functional cerebral vasculature. For example, a targeted deletion of miR-126 in mice showed that the cerebral hemorrhage due to defective vascular integrity accompanied a severe reduction in cranial vessel formation (Wang et al, 2008). In addition, excessive angiogenesis was shown to precede the cerebral hemorrhage in the developing brain of neuroendothelium-specific *itgb8* knockout mice (Arnold et al, 2014). Based on the mutant phenotypes described in our report, it is possible that DYRK1A may be a novel dual-function player that regulates both cerebral angiogenesis and the maintenance of vascular integrity simultaneously to prevent hemorrhage in the developing brain. Consistent with this idea, it has been reported that DS fetuses displayed several developmental vascular defects revealed by ultrasound scanning (Chaoui, 2005), while DS adults suffered a significantly higher risk of hemorrhagic stroke, as shown in a large cohort study (Sobey et al, 2015), suggesting potential dual roles of *DYRK1A* in these vascular disorders.

*Dyrk1aa* function in angiogenesis and hemorrhage prevention is likely to be a calcium-dependent process, based on the comparative transcriptome analyses of the WT and *dyrk1aa* mutants (Fig. 7). The Differentially-Enriched Gene (DEG) analysis identified “calcium ion binding” as the top-ranked GO category, which may reflect dysregulation of calcium homeostasis in *dyrk1aa* mutants. Of interest, several genes in the “calcium ion binding” category have been associated with endothelial cell permeability and vascular dysfunction. For example, Protocadherins such as *pcdhs-gamma*, one of the prominent DEGs from our analysis, were recently reported to be highly expressed in endothelial cells of the brain microvasculature and may contribute to junctional stability of the blood-brain barrier (Dilling et al, 2017). In addition, C2 Calcium Dependent Domain Containing 4A *(c2cd4a*, *zgc:162595*) is primarily expressed in endothelial cells, potentially increasing vascular permeability upon inflammation via transcriptional regulation related to cell adhesion (Warton et al, 2004). Furthermore, knockdown of the LDL receptor (*ldlr*) in zebrafish resulted in lipid accumulation in the blood and vascular leakage (O’Hare et al, 2014). Altered DEGs in the “oxidoreductase activity” MF category and “oxidation-reduction process” BP category in *dyrk1aa* mutants are also consistent with the disruption of Ca^2+^ signaling, because redox homeostasis is one of the major factors in regulating intracellular Ca^2+^ signaling events as well as vascular development (Lounsbury et al, 2000).

The calcium signal-related function of *dyrk1aa* in cerebrovascular formation and maintenance is also supported by results from the *in vivo* chemical screening and calcineurin inhibition studies. First, the specific calcium chelator EGTA was identified as one of the most potent small molecules that rescued the cerebral hemorrhage elicited by DYRK1A inhibition (Fig. 6). The Ca^2+^ ion level inside the cytosol is normally maintained at a low concentration (approximately 100 nM), compared to a significantly higher (more than 20,000-fold) concentration in the extracellular environment (Clapham, 2007), illustrating the importance of the tight regulation of calcium homeostasis across the membrane in maintaining normal cellular functions. As EGTA is an extracellular calcium chelator, it may affect the calcium function either extracellularly by directly attenuating the activities of calcium-dependent extracellular vascular ligands and cell adhesion molecules (Deli, 2009; Tomita et al, 1996) or intracellularly by lowering the amount of calcium entry into the cells. Our data suggested that both scenarios would be possible based on the facts that cerebral vessels in the mutants were disrupted at the ultrastructural level, presumably due to the altered calcium-dependent vascular integrity (Fig. 4), and rescue of the mutants’ vascular phenotypes were mimicked by inhibiting a calcium-dependent intracellular signaling pathway using FK506, a specific calcineurin inhibitor (Fig. 8). Consistent with these findings, dantrolene, a ryanodine receptor antagonist that decreases the intracellular calcium level (Fruen et al, 1997), was also identified as a modestly effective small molecule in our chemical screening, although it did not reach the statistical significance in our detailed analyses (data not shown).

An underlying mechanism by which *dyrk1aa* regulates the calcium signaling in vascular formation is not yet clear. DYRK1A may affect Ca^2+^ flux directly as in the case of GluN2A-containing NMDARs (N-methyl-D-aspartate glutamate receptors) phosphorylation by DYRK1A, leading to the elevation of their density on the membrane, and eventually on neurological dysfunction (Grau et al, 2014). Alternatively, Dyrk1a may indirectly influence Ca^2+^ signaling by phosphorylating mediators that determine the expression or activity of calcium-dependent effectors, similar to myocardial pathology, in which phosphorylation of Alternative Splicing Factor (ASF) by Dyrk1a increases the expression of Ca^2+^/calmodulin-dependent protein kinase II δ (CaMKIIδ)(He et al, 2015). Because Dyrk1a is likely to be functional both in the nucleus and in the cytoplasm based on its ubiquitous localization at the cellular level (Marti et al, 2003), it may directly or indirectly regulate membrane-bound ion channels near the cell membrane (similar to GluN2A phosphorylation), intracellular signaling molecules in the cytoplasm, or transcription factors in the nucleus, eventually maintaining the Ca^2+^ homeostasis of cells.

Together, our characterization of zebrafish *dyrk1aa* knockout mutants showed that *dyrk1aa* is implicated in cerebral angiogenic activity and the maintenance of vascular integrity during development. The combination of detailed transcriptomic analyses and chemical screening results strongly suggested that the calcium-dependent signaling regulated by *DYRK1A* is one of the major signaling pathways responsible for such vascular phenotypes. However, detailed signaling molecules and pathways affected in *dyrk1aa* mutants remain to be further studied. Our results also illustrate the usefulness of zebrafish *dyrk1aa* mutants in providing an *in vivo* animal model to understand the pathophysiology of human vascular diseases related to *DYRK1A* function, and suggest potential therapeutic approaches for effective treatments.

## MATERIALS AND METHODS

### Maintaining zebrafish embryos

Zebrafish (Danio rerio) embryos of AB (wild type) strain, transgenic zebrafish *Tg(kdrl:EGFP)* and *dyrk1aa* knockout mutant (*dyrk1aa*^*krb1*^) were maintained in E3 Egg water (5 mM NaCl, 0.17 mM KCl, 0.33 mM CaCl_2_, 0.33 mM MgSO_4_) in a petri dish at 28.5°C. In order to generate transparent zebrafish embryos for imaging confocal microscopy, performing whole mount i*n situ* hybridization, and examining hemorrhagic phenotype, embryos were incubated in 1X PTU (0.003% 1-phenyl 2-thiourea, Sigma-Aldrich)-E3 egg water after 6 hours post fertilization (hpf). Zebrafish husbandry and animal care were performed in accordance with guidelines from the Korea Research Institute of Bioscience and Biotechnology (KRIBB) and approved by KRIBB-IACUC (approval number: KRIBB-AEC-17126).

### *o*-dianisidine staining and quantification of the hemorrhagic phenotype

Embryos at 52 hpf were fixed with 4% paraformaldehyde in 1X phosphate-buffered saline (PBS) for 4 hours at room temperature (RT) and washed with 1X PBS containing 0.1% Tween 20 (1X PBST). The embryos were placed in *o*-dianisidine stain solution (0.6 mg/ml *o*-dianisidine (Sigma), 0.01M sodium acetate (pH 5.5), 0.65% hydrogen peroxide and 40% ethanol) in the dark for several minutes at RT to detect hemoglobin activity. *o*-dianisidine is a peroxidase substrate, and hemoglobin catalyzes the H_2_O_2_ mediated oxidation of *o*-dianisidine. Stained embryos were washed several times with 1X PBST and stored in 70% glycerol for imaging using an Olympus SZX16 microscope equipped with TUCSEN Dhyana 400DC. The cerebral hemorrhagic phenotype was calculated as the mean percentages by counting the numbers of embryo with hemorrhagic phenotype in the brain and the retina.

### Fluorescence-Assisted Cell Sorting (FACS) analysis of endothelial cells

To isolate GFP positive endothelial cells from *Tg(kdrl:EGFP)* embryos, we adopted the protocol developed by Manoli et al.(Manoli & Driever, 2012) Briefly, 150 embryos of at 52 hpf were dechorinated with 1 mg/ml protease (P6911, Sigma) in E3 Egg water for 5 min at RT, and washed in 0.5 × Danieau’s solution (29 mM NaCl, 0.35 mM KCl, 0.2 mM MgSO_4_•7H_2_O, 0.3 mM Ca(NO_3_)_2_, 2.5 mM HEPES, pH 7.6). The yolk of those embryos were removed with the deyolking buffer (55 mM NaCl, 1.8 mM KCl, 1.25 mM NaHCO_3_), followed by washing in 0.5X Danieau’s solution, and embryonic cells were dissociated by squishing the cell suspension with a 40 μm strainer (93040, SPL) using FACS max cell dissociation solution (AMS Biotechnology, Abingdon, UK; T200100). FACS was performed at RT under sterile conditions using a FACSAria (BD Biosciences).

### RNA preparation and RT-PCR analysis

Zebrafish embryos in each developmental stage or isolated cells by FACS were harvested with TRI reagent solution (Ambion, USA), followed by purifying total RNA with Direct-zol RNA miniprep kit (Zymo Research) and synthesizing cDNA with SuperScript III First-Strand Synthesis System (Invitrogen, USA). The synthesized cDNA was amplified by PCR using the forward primer 5’-TCA GTG ATG CTC ACC CAC AG-3’ and reverse primer 5’-CGT CAT AGC CGT CGT TGT AA-3’ for *dyrk1aa*, the forward primer 5’- GGC AAG CTG ACC CTG AAG TT-3’ and reverse primer 5’- TTC TGC TTG TCG GCC ATG AT-3’ for *EGFP*, the forward primer 5’- CCC TTA CCC TGG CTT ACA CA-3’ and reverse primer 5’- TCT TGT TGG TTC CGT TCT CC-3’ for *kdr1*, the forward primer 5’- CTG GTT CAA GGG ATG GAA GA-3’ and reverse primer 5’- ATG TGA GCA GTG TGG CAA TC-3’ for *eef1a1l1*.

### Chemical treatment and small molecule library screening

Harmine (Sigma) was dissolved in DMSO and added to 1X PTU-E3 Egg water to earn final concentrations of 10-50 μM with 0.1% DMSO, and then applied to approximately 40 embryos (dechorionated) of 90 mm plate from 24 to 52 hpf.

For chemical screening, each of 1280 small molecules in the Library of Pharmacologically Active Compounds (LOPAC1280, Sigma) with 10 μM as a final concentration was individually applied into each well of 48 well plates containing 1X PTU-E3 Egg water with 30 μM harmine and five embryos from 24 to 52 hpf. As a negative control, 1XPTU-E3 Egg water containing 0.4% DMSO was used. Upon 30 μM harmine treatment, three embryos on average displayed the hemorrhage in the brain regions. Based on this criteria, the increased hemorrhage was defined by 4 or 5 embryos showing the hemorrhage in brain regions, whereas the reduced phenotype was defined by 0 to 2 embryos with such defect.

For EGTA and FK506 treatment, EGTA at final concentrations of 1-100 nM and FK506 at final concentrations of 10-100 ng/ml was treated into each well of 6-well plates containing approximately 20 dechorionated embryos of WT and *dyrk1aa* mutants from 24 to 52 hpf, with 1XPTU-E3 Egg water containing 0.1% DMSO alone used as a negative control.

### Confocal microscopic analysis for cerebrovascular phenotypes of zebrafish embryos

To analyze the brain vasculature phenotypes of *Tg(kdrl:EGFP)* embryos with high resolution, embryos were grown up to 52 hpf and fixed with 1X staining solution (4% paraformaldehyde, 4% sucrose, 0.15 mM CaCl_2_, 1X PBS) overnight at 4°C. Fixed embryos were washed briefly with 1X PBST and embedded on the glass-bottomed imaging dishes with 1% LMPA (low melting point agarose, Promega). The embryos were imaged using Olympus FV1000 confocal microscopy and the central artery (CtA) development in the hindbrain was quantified with the length and branching points by measuring total lengths with Image J and manually counting the junctional site of the CtAs.

### Whole mount in situ hybridization and section of the hybridized embryos

Whole-mount *in situ* hybridization (WISH) in zebrafish embryos was performed as previously reported(Thisse & Thisse, 2008). The DNA templates for zebrafish *dyrk1aa* and *dll4* (GeneBank accession numbers BC129212.1 and NM_001079835.1, respectively) were amplified from cDNA of WT embryos at 52 hpf. The DNA of *kdrl*, *krox20* and *isl1* were donated from Porf. Cheol-Hee Kim of Chungnam National University. Dig-labelled antisense probes were *in vitro* transcribed by SP6 or T7 RNA polymerase kit (Roche) and purified with NucAway spin columns (Invitrogen). Embryos for WISH were prepared by fixing with 4% paraformaldehyde in 1X PBS solution, dehydrating using methanol, stored at −20°C over 30 minutes, and serially rehydrated with 1X PBST solution. The rehydrated embryos were treated with proteinase K in 1X PBS and post-fixed with 4% paraformaldehyde. The antisense probes were hybridized with the fixed embryos at each developmental stage in hybridizing solution (5 mg/ml torula yeast RNA type VI, 50 μg/ml heparin, 50% formamide, 5X SSC, 0.1X Tween-20, 1M citric acid used to adjust pH 6.0) at 70°C overnight. The probes were washed serially using 2X SSCT-F (2X SSCT, 50% formamide, 0.1% Tween20), 2X SSCT (2X SSCT, 0.1% Tween20), 0.2X SSCT (0.2X SSCT, 0.1% Tween20) at 70°C and 1X PBST at RT. The embryos were blocked with the blocking solution (5% horse serum, 1X PBST) at RT, and the alkaline phosphatase conjugated anti-digoxigenin antibody (Roche, 11 093 274 910) was added into the blocking solution at 4°C overnight. To detect the expression signal of transcript, NBT/BCIP solution (Roche, 11 681 451 001) was used as alkaline phosphatase substrate. The expression pattern of transcripts was observed by using Olympus SZX16 microscope and imaged with TUCSEN Dhyana 400DC or Olympus XC10. To observe detailed expression pattern of *dyrk1aa*, in situ hybridized embryos were prepared for cryosectioning by embedding in an agar-sucrose solution (1.5% agar, 5% sucrose). After the agar blocks containing the embryos were kept in 30% sucrose solution, they were processed for transverse cryosectioning with a thickness of 25-35 μm.

### Microinjection of RNA

In order to prepare mRNAs of *dyrk1aa*, *dyrk1aa-K193R* and *mCherryRed* for rescue experiments, the PCS2+ vectors inserted with each DNA were linearize, *in vitro* transcribed by mMESSAGE mMACHINE kit (Invitrogen), and purified with NucAway spin columns (Invitrogen). 1-cell stage eggs were collected and microinjected with each mRNA construct containing 1% phenol red solution as a visible indicator using PV380 Pneumatic picopump (World Precision Instruments).

### Transmission electron microscopy

Tissue samples from embryos of WT and *dyrk1aa*^*krb1*^ at 48 hpf were fixed immediately with 2% glutaraldehyde and 2% paraformaldehyde in 0.1 M phosphate buffer (pH 7.4) for 2 h at 4°C. Following three washes in the phosphate buffer, tissues were post-fixed with 1% osmium tetroxide on ice for 2 hours and washed three times, all in the phosphate buffer. The tissues were then embedded in pure Epon 812 mixture after dehydration in ethanol series and followed by infiltration in propylene oxide:epon mixture series. Polymerization was conducted with pure resin at 70°C for 24 hours. Ultrathin sections (~70 nm) were obtained with a model MT-X ultramicrotome (RMC, Tucson, AZ,) and then collected on 100 mesh copper grids. After staining with 2% uranyl acetate (7 min) and lead citrate (2 min), the sections were visualized by Bio-HVEM system (JEM-1400Plus at 120 kV and JEM-1000BEF at 1000 kV, JEOL, JAPAN).

### Isolation, Library preparation and sequencing for RNA seq

Total RNA was isolated using Trizol reagent (Invitrogen). RNA quality was assessed by Agilent 2100 bioanalyzer using the RNA 6000 Nano Chip (Agilent Technologies, Amstelveen, The Netherlands), and RNA quantification was performed using ND-2000 Spectrophotometer (Thermo Inc., DE, USA). For control and test RNAs, the construction of library was performed using SENSE mRNA-Seq Library Prep Kit (Lexogen, Inc., Austria) according to the manufacturer’s instructions. Briefly, each 2 ug total RNA are prepared and incubated with magnetic beads decorated with oligo-dT and other RNAs except mRNA was removed by washing solution. Library production was initiated by the random hybridization of starter/stopper heterodimers containing Illumina-compatible linker sequences to the poly (A) RNA bound to the magnetic beads. A single-tube reverse transcription and ligation reaction extends the starter to the next hybridized heterodimer, where the newly-synthesized cDNA insert is ligated to the stopper. Second strand synthesis was performed to release the library from the beads, and the library was then amplified. Barcodes were introduced when the library is amplified. High-throughput sequencing was performed as paired-end 100 sequencing using HiSeq 2500 (Illumina, Inc., USA). The sequenced reads were mapped to the UCSC zebrafish genome (danRer10) using STAR (v.2.5.1) (Dobin et al., 2013), and the gene expression levels were quantified with the count module in STAR. The edgeR (v.3.12.1) (Robinson et al., 2010) package was used to select differentially expressed genes from the RNA-seq count data. Meanwhile, the TMM (trimmed mean of M-values)-normalized CPM (counts per million) value of each gene was set to a baseline of 1 and log2-transformed for further analysis (i.e., volcano plot drawing and correlation analysis). The results of RNA-seq data were deposited in NCBI (GEO: GSE111280).

### Statistical analyses

Statistical analyses of the data were performed with the student’s *t*-test or one-way ANONA with Dunnett’s multiple comparisons test using Prism software (Ver.7). Data are shown as mean ± S.E.M. with *p*-values (\**p* < 0.05, \*\**p* < 0.01, \*\*\**p* < 0.005).

## ACKNOWLEDGMENTS

This work was supported by National Research Foundation (NRF) and National Research Council of Science & Technology (NST) grant of the Korea government (MSIP) (NRF-2011-0023507, CRC-15-04-KIST), and KRIBB Research Initiative Program.

## THE PAPER EXPLAINED

PROBLEM: *DYRK1A* (*Dual-specificity tyrosine phosphorylation regulated kinase 1A*) is a major causative gene in Down syndrome (DS), and this gene is related to various DS symptoms. Since DS patients displayed abnormal vasculatures and the reduction in solid tumor incidence requiring the formation of new vasculatures, we questioned whether the *DYRK1A* gene has function in vascular formation.

RESULTS: To identify *DYRK1A* gene function of vascular formation, we used zebrafish *dyrk1aa* mutant embryos, which exhibited bleeding and abnormal vascular development in the brain. By performing a chemical screening, we found that a calcium chelator, ethylene glycol-bis(β-aminoethyl ether)-N,N,N’,N’-tetraacetic acid (EGTA) was able to rescue the bleeding and abnormal vascular development in zebrafish *dyrk1aa* mutants. We also showed that FK506, another small molecule that modulates calcium signaling within a cell, was able to rescue these vascular defects. In addition, the whole gene expression comparison confirmed the alteration of calcium signaling networks in *dyrk1aa* mutants, further underscoring the importance of calcium signaling in the vascular defects of *dyrk1aa* mutants.

IMPACT: We found essential, but under-explored, roles of *dyrk1aa* in vascular development via regulation of calcium signaling in the living organism. These novel findings may apply for the *DYRK1A*-related vascular disease therapy.

## AUTHOR CONTRIBUTIONS

HJC, JGL, JHK, YHH, HJK, SYK, and JSL performed experiments and analyzed the data. KSL and KY helped data interpretation and discussion. HJC and JSL designed the study and wrote the manuscript.

